# Evaluating the efficacy of Optoα1AR activation in astrocytes in modulating basal hippocampal synaptic excitation and inhibition

**DOI:** 10.1101/2021.01.06.425606

**Authors:** Connor D. Courtney, Courtney Sobieski, Charu Ramakrishnan, Robbie J. Ingram, Natalia M. Wojnowski, R. Anthony DeFazio, Karl Deisseroth, Catherine A. Christian-Hinman

**Affiliations:** Neuroscience Program, University of Illinois at Urbana-Champaign, Urbana, IL 61801; Department of Molecular and Integrative Physiology, University of Illinois at Urbana-Champaign, Urbana, IL 61801; Beckman Institute for Advanced Science and Technology, University of Illinois at Urbana-Champaign, Urbana, IL 61801; Department of Bioengineering, Stanford University, Stanford, CA 94305; Molecular and Integrative Physiology, University of Michigan, Ann Arbor, MI 48109

## Abstract

Astrocytes play active roles at synapses and can monitor, respond, and adapt to local synaptic activity. To investigate this relationship, more tools that can selectively activate native G protein signaling pathways in astrocytes with both spatial and temporal precision are needed. Here, we tested AAV8-GFAP-Optoα1AR-eYFP (Optoα1AR), a viral vector to enable activation of G_q_ signaling in astrocytes via light-sensitive α1-adrenergic receptors. To determine if stimulating astrocytic Optoα1AR modulates hippocampal synaptic transmission, recordings were made in CA1 pyramidal cells with surrounding astrocytes expressing Optoα1AR, channelrhodopsin (ChR2), or GFP. Both high-frequency (20 Hz, 45-ms light pulses, 5 mW, 5 min) and low-frequency (0.5 Hz, 1-s pulses at increasing 1, 5, and 10 mW intensities, 90 s per intensity) blue light stimulation were tested. 20 Hz Optoα1AR stimulation increased both inhibitory and excitatory postsynaptic current (IPSC and EPSC) frequency, and the mIPSC effect was largely reversible within 20 min. By contrast, low-frequency stimulation of Optoα1AR did not modulate either IPSCs or EPSCs, whereas the same stimulation of astrocytic ChR2 was effective. These data demonstrate that Optoα1AR activation in astrocytes changes synaptic excitation and inhibition in a stimulation-sensitive manner, demonstrating the efficacy and utility of GFAP-Optoα1AR as a tool in studying astrocyte-neuron interactions.

## Introduction

Astrocytes can detect synaptic signaling through G protein-coupled receptor (GPCR) activation^1, 2^ and respond to local neuronal activity with transient increases in intracellular Ca^2+^ levels^3, 4^. As influential components of the tripartite synapse, astrocytes engage in extensive and specific bidirectional communication with synapses^5–7^. In the hippocampus, many studies investigating astrocyte-neuron interactions have demonstrated astrocyte-specific modulation of excitatory transmission and/or plasticity^8–16^. Evidence for astrocytic regulation of basal synaptic inhibition remains limited, although previous studies have suggested a role for astrocyte-mediated modulation of fast synaptic inhibition in multiple brain areas including hippocampus, cortex, and thalamus^17, 18^. For example, mechanical stimulation of astrocytes leads to glutamate release and a strengthening of inhibition that is dependent upon astrocytic Ca^2+^ signaling and AMPA/NMDA receptors^19^. In addition, activation of somatostatin-positive interneurons stimulates the release of ATP/adenosine from astrocytes and subsequent enhancement of inhibition via A_1_ adenosine receptors^20^, suggesting that astrocytes can modulate both inhibitory and excitatory synaptic transmission.

Historically, the study of neuron-astrocyte interactions has been hampered by the limited availability of tools that selectively modulate astrocytic GPCR signaling with precise spatial and temporal control. Several studies have utilized Designer Receptors Exclusively Activated by Designer Drugs (DREADDs) for specific manipulation of astrocytes^8, 21–24^. DREADDs are well-suited for this purpose as they provide targeted activation of GPCR-mediated endogenous intracellular cascades native to astrocytes. A limitation of DREADDs, however, is the relative lack of temporal regulation of the activation of GPCR-mediated signaling. To overcome this drawback, in many studies the light-gated nonspecific cation channel channelrhodopsin-2 (ChR2) has been inserted into astrocyte membranes for improved temporal control^25–31^. ChR2 activation, however, produces robust depolarization that likely exceeds that endogenously produced in astrocytes, and increases extracellular K^+^ levels^30^. Furthermore, ChR2 activation does not recapitulate a physiologically relevant signaling cascade to drive intracellular Ca^2+^ elevations in astrocytes^32^. Therefore, there remains a need for optical tools that allow for precise spatial and temporal stimulation of astrocytes with improved physiological relevance. Progress has been made in this regard with the recent development of an astrocyte-specific melanopsin coupled to the G_q_ intracellular pathway^9^. The use of OptoXRs^33^ has emerged as another potential solution, and stimulation of Optoα1 adrenergic receptor (Optoα1AR)-derived constructs can drive Ca^2+^ elevations in cultured astrocytes^34^ and stimulate memory enhancement *in vivo*^8^.

Here, we generated a novel adeno-associated virus (AAV) construct, AAV8-GFAP-Optoα1AR- eYFP, to efficiently insert Optoα1AR specifically into astrocytes. We report that stimulation of astrocytic Optoα1AR modulates basal synaptic transmission in CA1, and this response depends on the properties of the light stimulation. These data suggest that this vector may be a useful tool in the studies of astrocytic modulation of neuronal and synaptic function.

## Results

### Astrocyte-specific expression of optogenetic constructs

AAV8 vectors driven by the astrocyte-specific GFAP promoter were individually injected bilaterally into dorsal CA1 of wild-type C57BL/6J mice. For each mouse, one of three constructs was used: GFAP-Optoα1AR-eYFP (Optoα1AR), GFAP-hChR2(H134R)-eYFP (ChR2), and GFAP-GFP (GFP). To confirm the astrocyte-specificity of these vectors, immunohistochemistry was performed for the neuronal marker NeuN. Colocalization was not observed between NeuN staining of CA1 pyramidal cells and the GFP or eYFP fluorescent tags of any AAV used (**Figure 1**), indicating a lack of neuronal expression of the optogenetic constructs.

**Figure 1.**
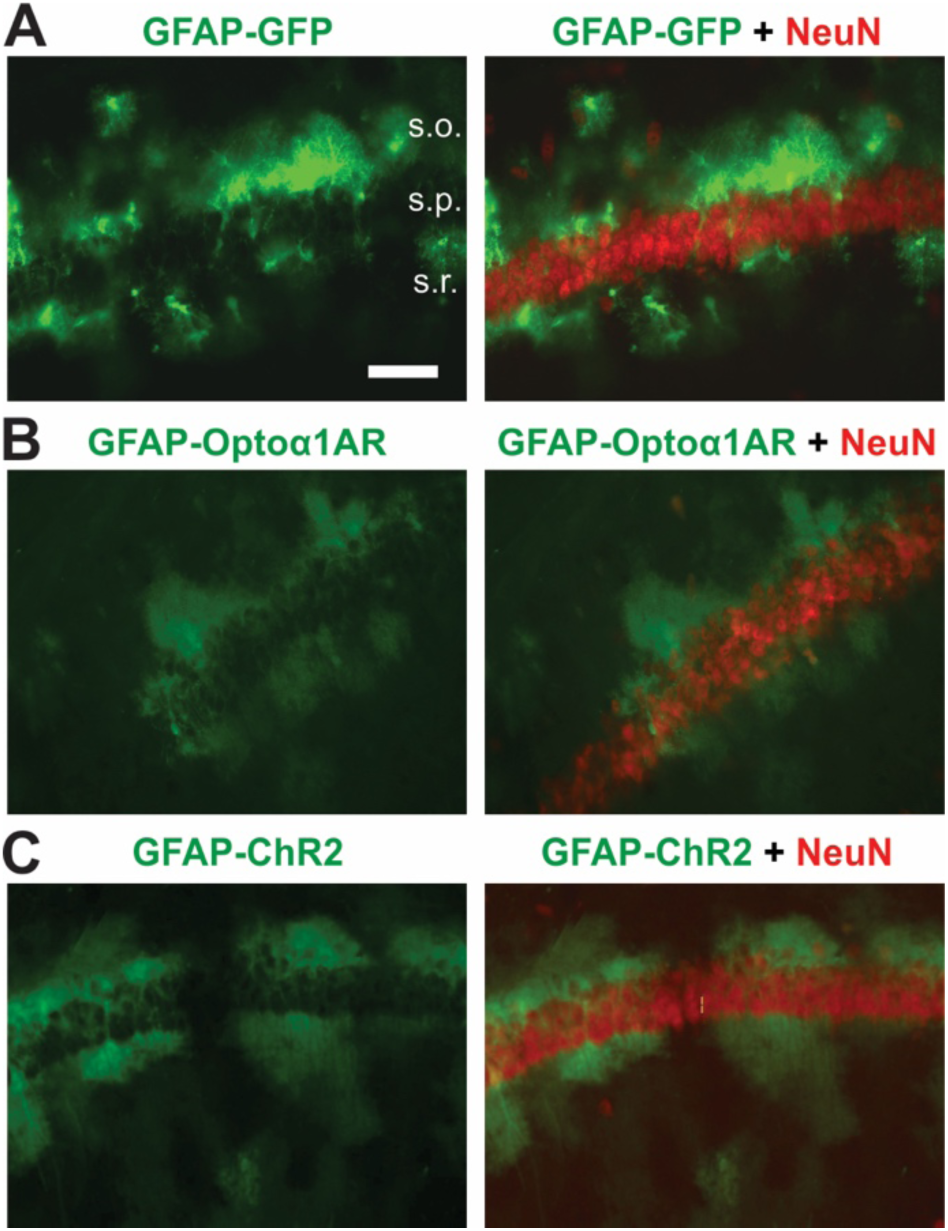
AAV8-GFAP-Optoα1AR-eYFP selectively targets astrocytes in hippocampal CA1. Example immunohistochemical images displaying astrocytic targeting of optogenetic constructs. (A) AAV8-GFAP-GFP (left) and AAV8-GFAP-GFP merged with neuronal marker NeuN (right). s.o., stratum oriens; s.p., stratum pyramidale; s.r., stratum radiatum. Scale bar, 40 μm (B) AAV8-GFAP-Optoα1AR-eYFP (left) and AAV8-GFAP-Optoα1AR-eYFP merged with NeuN (right). (C) AAV8-GFAP-hChR2(H134R)-eYFP (left), and AAV8-GFAP-hChR2(H134R)-eYFP merged with NeuN (right).

### High-frequency (20 Hz) optogenetic stimulation of Optoα1AR in astrocytes produces a sustained modulation of sIPSC frequency

High-frequency optical stimulation of astrocytes using G_q_-coupled melanopsin enhances inhibitory transmission in the medial prefrontal cortex^35^, and 20 Hz stimulation of a similar OptoG_q_ construct in CA1 astrocytes improves memory performance *in vivo*^8^. To examine if 20 Hz activation of Optoα1AR in astrocytes modulates basal synaptic inhibition in hippocampus, whole-cell patch clamp recordings of spontaneous inhibitory postsynaptic currents (sIPSCs) were made in CA1 pyramidal cells. Slices containing astrocytes expressing either Optoα1AR or control GFP were exposed to 20 Hz blue light stimulation at 5 mW intensity for 5 minutes (closely resembling *in vivo* experiments described in ref. 8) (**Figure 2A**). First, we investigated the impact of 20 Hz activation of Optoα1AR or GFP in astrocytes in modulating hippocampal inhibitory transmission. 20 Hz stimulation of the Optoα1AR group led to an overall increase in sIPSC frequency compared to baseline when the analysis incorporated all sIPSCs recorded during the duration of light stimulation together (t=-2.73, p=0.015, n=16 cells from 5 mice, 9 of 16 cells displaying >20% increase), and this effect was not seen in the control GFP group (t=0.67, p=0.51, n=8 cells from 2 mice) (**Figure 2B, 2C**). To investigate the time course of the effect, sIPSC frequency was subdivided into 30-s bins. This analysis, however, did not yield a significant difference in sIPSC frequency for any 30-s bin compared to baseline in either the Optoα1AR group (p=0.40) or the GFP group (p=0.86) (**Figure 2D**), reflecting the variability in time course of the effect within individual cells. In addition, sIPSCs from the Optoα1AR group displayed no overall change in amplitude during 20 Hz stimulation compared to baseline (t=0.18, p=0.86) (**Figure 2E**). When subdivided into 30-s intervals, however, sIPSC amplitude reached its average maximum at 60-90 s into the light stimulation, and its average minimum at 240-300 s, with significant differences in amplitude between these maximum and minimum timepoints (p=0.02) (**Figure 2F, top**). By contrast, GFP controls did not demonstrate any changes in sIPSC amplitude either overall (t=-0.01, p=0.99) or across time (p=0.86) (**Figure 2E; 2F, bottom**).

**Figure 2.**
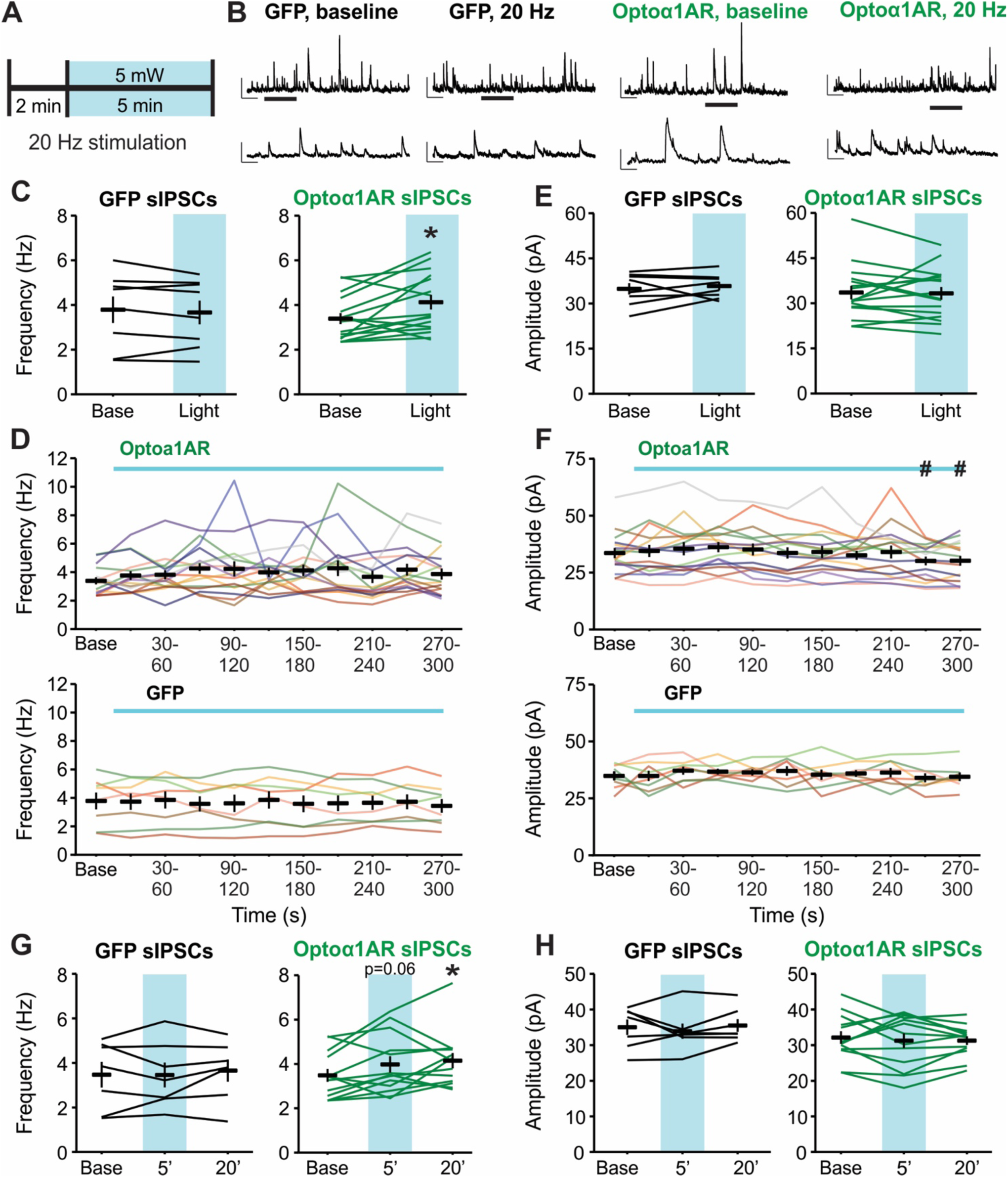
High-frequency (20 Hz) stimulation of astrocytic Optoα1AR increases sIPSC frequency. (A) Schematic representation of 20 Hz stimulation paradigm. (B) Representative sIPSC traces from individual pyramidal cells before (left) and during (right) 20 Hz stimulation of GFP (left two traces) or Optoα1AR (right two traces) in astrocytes. Insets represent expanded sections from underlined portion of trace. Scale bars: 50 pA, 1 s; inset: 50 pA, 200 ms. (C) Mean ± SEM of sIPSC frequency before (Base) and during the full duration of 20 Hz stimulation (Light) of GFP (left) or Optoα1AR (right) in astrocytes. Lines represent individual cells. (D) Mean ± SEM of sIPSC frequency across time of Optoα1AR (top) and GFP (bottom) during 20 Hz stimulation. Lines represent individual cells. Note that lines are used to enable identification of the same cells across the time points; increased slope at end of baseline period does not represent an increase in frequency prior to light delivery. (E) Mean ± SEM of sIPSC amplitude before (Base) and during the full duration of 20 Hz stimulation (Light) of GFP (left) or Optoα1AR (right) in astrocytes. Lines represent individual cells. (F) Mean ± SEM of sIPSC amplitude across time in the Optoα1AR (top) and GFP (bottom) groups during 20 Hz stimulation. Lines represent individual cells. Note that lines are used to enable identification of the same cells across the time points; increased slope at end of baseline period does not represent an increase in frequency prior to light delivery. (G) Mean ± SEM of sIPSC frequency before (Base), during the final minute of 20 Hz stimulation (5’), and 20 minutes following 20 Hz stimulation (20’) of GFP (left) or Optoα1AR (right) in astrocytes. Lines represent individual cells. (H) Mean ± SEM of sIPSC amplitude before (Base), during the final minute of 20 Hz stimulation (5’), and 20 minutes following 20 Hz stimulation (20’) of GFP (left) or Optoα1AR (right) in astrocytes. Lines represent individual cells. *, p<0.05 vs. baseline value. #, p<0.05 vs. 60-90 s timepoint.

An important advantage of using optogenetic tools is the temporal precision of activation^36^, and Optoα1AR expression in neurons allows for spatiotemporally precise manipulation of biochemical signaling and modulation of behavior *in vivo*^33^. In theory, this temporal control could allow for cellular modulation to be reversible at short timescales. To test for this possibility, we recorded sIPSCs for 20 additional min following cessation of the 20 Hz light stimulation in a subset of cells in slices expressing Optoα1AR or GFP in astrocytes. In these cells, 20 Hz stimulation of Optoα1AR resulted in a trend for increased sIPSC frequency during the final minute of stimulation compared to baseline (p=0.06, n=13 cells), similar to the modulation of sIPSC frequency observed in the full group (**Figure 2C**). Contrary to our prediction, however, cells from the Optoα1AR group maintained an increased level of sIPSC frequency at 20 min post-stimulation (p=0.02 compared to baseline, 8 of 13 cells displaying >20% increase) (**Figure 2G**). This finding suggests that 20 Hz stimulation of Optoα1AR in astrocytes can exert long-lasting modulation of synaptic inhibition. In cells from the GFP group, by contrast, no changes in sIPSC frequency were seen either during the final minute of 20 Hz stimulation or at 20 min post-stimulation (n=7 cells from 2 mice) (**Figure 2G**). Additionally, there were no significant changes in sIPSC amplitude either during the light stimulation or at 20 min afterwards in cells from the Optoα1AR or GFP groups (**Figure 2H**). Altogether, these data suggest that high-frequency stimulation of Optoα1AR in astrocytes can modulate hippocampal synaptic inhibition, and that this modulation is sustained for at least 20 minutes following termination of the stimulation.

### High-frequency (20 Hz) optogenetic stimulation of Optoα1AR in astrocytes enhances activity-independent hippocampal synaptic inhibition

To determine if the effects of 20 Hz stimulation of astrocytic Optoα1AR on hippocampal synaptic inhibition require presynaptic input, miniature inhibitory postsynaptic currents (mIPSCs) were recorded under the same stimulation paradigm in the presence of tetrodotoxin (TTX, 0.5 μM) to prevent action potential firing (**Figure 3A**). mIPSCs recorded from the Optoα1AR group displayed an increase in frequency in response to 5 min of 20 Hz light stimulation (t=-3.91, p=0.002, n=14 cells from 6 mice, 10 of 14 cells displaying >20% increase) (**Figure 3B**). The increase in frequency was first visible after 2 minutes of light stimulation (p=0.01) and peaked within 4 minutes of stimulation (p<0.001) (**Figure 3C, top**). No increase in mIPSC frequency was seen in the control GFP group (t=-0.84, p=0.42, n=13 cells from 5 mice) (**Figure 3B**), suggesting that 20 Hz stimulation of Optoα1AR can acutely elicit an activity-independent mechanism producing increased frequency of GABA release onto CA1 pyramidal cells. mIPSC amplitude was not affected by light stimulation in the Optoα1AR group (t=-0.98, p=0.34) (**Figure 3D**). Additionally, no differences in mIPSC amplitude were observed over time (**Figure 3E, top**). However, there was an increase in mIPSC amplitude in cells from the GFP group (t=-4.10, p<0.001) (**Figure 3D**) that appeared within the first 30 s of light stimulation (**Figure 3E, bottom**). This effect may reflect a non-specific artifact of exciting GFP in astrocytes in the absence of an opsin, underscoring the necessity for fluorophore-only controls in optogenetic experiments. It should be noted, however, that this effect reflected a very small (∼2 pA) average increase, and the mean amplitude of events during the light stimulation was similar between groups.

**Figure 3.**
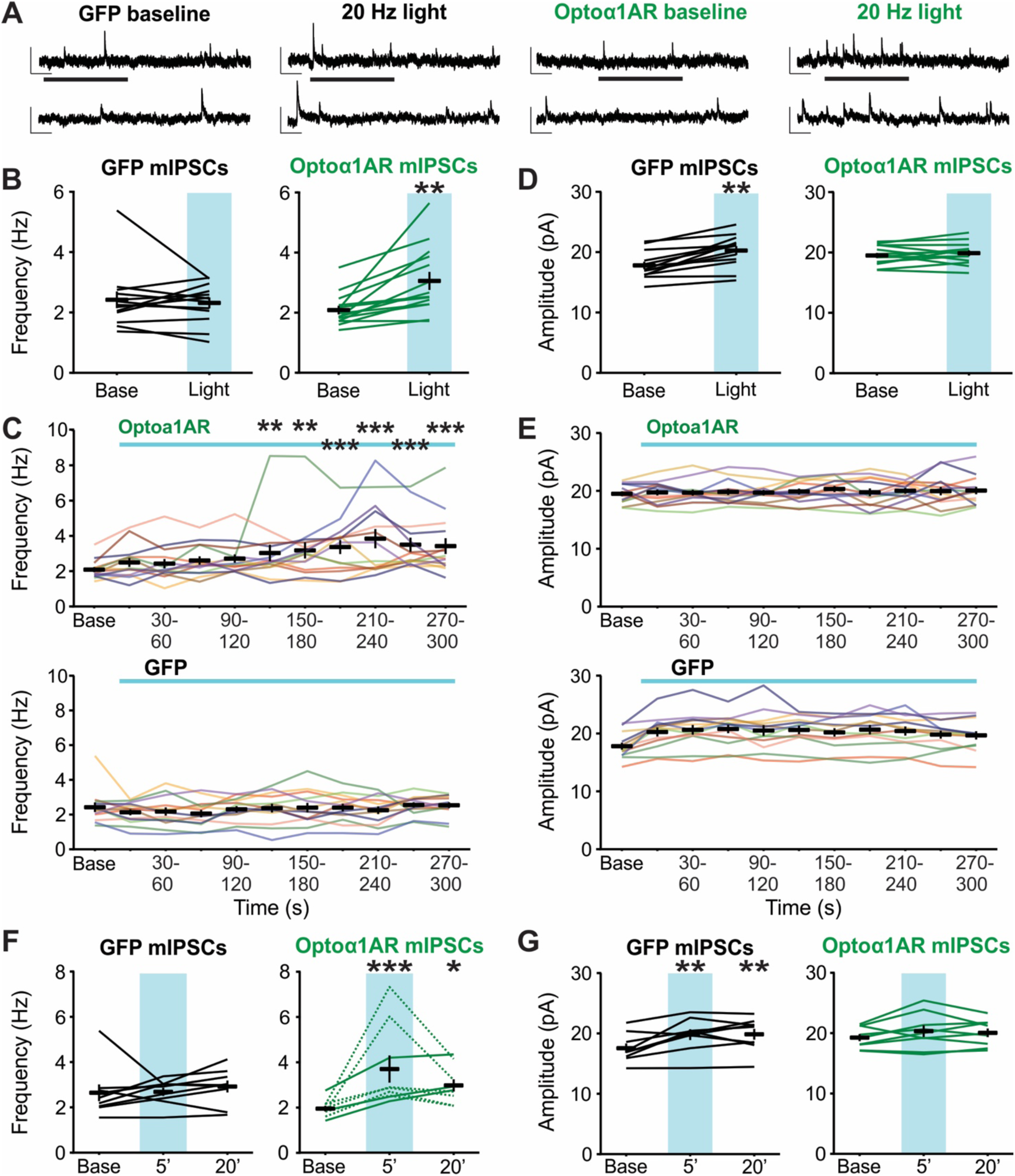
High-frequency (20 Hz) stimulation of astrocytic Optoα1AR increases mIPSC frequency. (A) Representative mIPSC traces from individual pyramidal cells before (left) and during (right) 20 Hz stimulation of GFP (left two traces) or Optoα1AR (right two traces) in astrocytes. Insets represent expanded sections from underlined portion of trace. Scale bars: 20 pA, 500 ms; inset: 20 pA, 100 ms. (B) Mean ± SEM of mIPSC frequency before (Base) and during the full duration of 20 Hz stimulation (Light) of GFP (left) or Optoα1AR (right) in astrocytes. Lines represent individual cells. (C) Mean ± SEM of mIPSC frequency across time in the Optoα1AR (top) and GFP (bottom) groups during 20 Hz stimulation. Lines represent individual cells. Note that lines are used to enable identification of the same cells across the time points; increased slope at end of baseline period does not represent an increase in frequency prior to light delivery. (D) Mean ± SEM of mIPSC amplitude before (Base) and during the full duration of 20 Hz stimulation (Light) of GFP (left) or Optoα1AR (right) in astrocytes. Lines represent individual cells. (E) Mean ± SEM of mIPSC amplitude across time in the Optoα1AR (top) and GFP (bottom) groups during 20 Hz stimulation. Lines represent individual cells. Note that lines are used to enable identification of the same cells across the time points; increased slope at end of baseline period does not represent an increase in frequency prior to light delivery. (F) Mean ± SEM of mIPSC frequency before (Base), during the final minute of 20 Hz stimulation (5’), and 20 minutes following 20 Hz stimulation (20’) of GFP (left) or Optoα1AR (right) in astrocytes. Lines represent individual cells. Dotted lines represent 6 cells that showed reversibility of effect. (G) Mean ± SEM of mIPSC amplitude before (Base), during the final minute of 20 Hz stimulation (5’), and 20 minutes following 20 Hz stimulation (20’) of GFP (left) or Optoα1AR (right) in astrocytes. Lines represent individual cells. *, **, ***, p<0.5, p<0.01, p<0.001 vs. baseline value

To determine if 20 Hz stimulation of astrocytic Optoα1AR leads to a sustained increase in mIPSC frequency, a subset of cells from the Optoα1AR and GFP groups were recorded for 20 additional min following cessation of the light stimulation. In cells from the Optoα1AR group, an increase in mIPSC frequency was observed during the last minute of 20 Hz light stimulation (p<0.001, 9 of 9 cells displaying >30% increase compared to baseline), and at 20 min following cessation of the stimulation (p=0.03, 7 of 9 cells displaying >20% increase compared to baseline) (**Figure 3F**). However, in contrast to the sIPSCs, 6 of 9 cells displayed a reduction in frequency at 20 minutes post-stimulation compared to the last minute of stimulation, and this reduction was greater than 50% in 5 of these 6 cells (**Figure 3F**). These data suggest that while 20 Hz stimulation of Optoα1AR in astrocytes leads to a sustained increase in mIPSC frequency, this activity-independent modulation is reversible to an extent that is not seen when firing activity is unblocked. In cells from the GFP group, no differences were seen in mIPSC frequency during light stimulation nor at 20 min after (9 cells from 4 mice) (**Figure 3F**). Additionally, there was no effect of 20 Hz stimulation on mIPSC amplitude in the Optoα1AR group at either timepoint. However, as observed in the full GFP mIPSC group, there was a small increase in mIPSC amplitude in the subgroup of cells recorded after light stimulation (p<0.01), and this increase persisted for at least 20 min (p<0.01). (**Figure 3G**). Altogether, these data indicate that high-frequency stimulation of Optoα1AR in astrocytes modulates activity-independent synaptic inhibition. In addition, although this stimulation drives an increase in mIPSC frequency that is both acute and sustained, this change appears more reversible than that observed when action potential firing is intact.

### High-frequency (20 Hz) optogenetic stimulation of Optoα1AR in astrocytes modulates activity-independent hippocampal glutamatergic transmission

Previous studies using G_q_-coupled optogenetic or chemogenetic tools have demonstrated that activation of this pathway in astrocytes can modulate basal excitatory neurotransmission in the hippocampus^8, 9^. As we had observed differences in IPSC frequency in response to 20 Hz stimulation of Optoα1AR in astrocytes, we sought to determine whether the same stimulation modulates mEPSCs recorded from CA1 pyramidal cells (**Figure 4A**). In the Optoα1AR group, this stimulation produced an increase in mEPSC frequency compared to baseline (t=-5.04, p<0.001, n=10 cells from 4 mice) (**Figure 4B**). The increase in frequency was first visible after 90 s of light stimulation (p=0.005) and was within 0.12 Hz of peak frequency at this time (90 s mean=2.77 ± 0.46 Hz; peak mean=2.89 ± 0.33 Hz) (**Figure 4C, top**), suggesting that activation of Optoα1AR in astrocytes may modulate glutamatergic transmission on a faster timescale compared to the effects on GABAergic transmission. No change in mIPSC frequency was observed in the control GFP group (t=0.13, p=0.91, n=6 cells from 4 mice) (**Figure 4B, 4C bottom**). Additionally, mEPSC amplitude was not changed in the Optoα1AR group either as compared to baseline (t=-0.56, p=0.59) (**Figure 4D**) or across time (**Figure 4E, top**), and no difference in amplitude was observed in the GFP group (t=-0.63, p=0.56) (**Figure 4D, 4E**). These data demonstrate that 20 Hz stimulation of Optoα1AR in astrocytes is capable of modulating hippocampal glutamatergic transmission.

**Figure 4.**
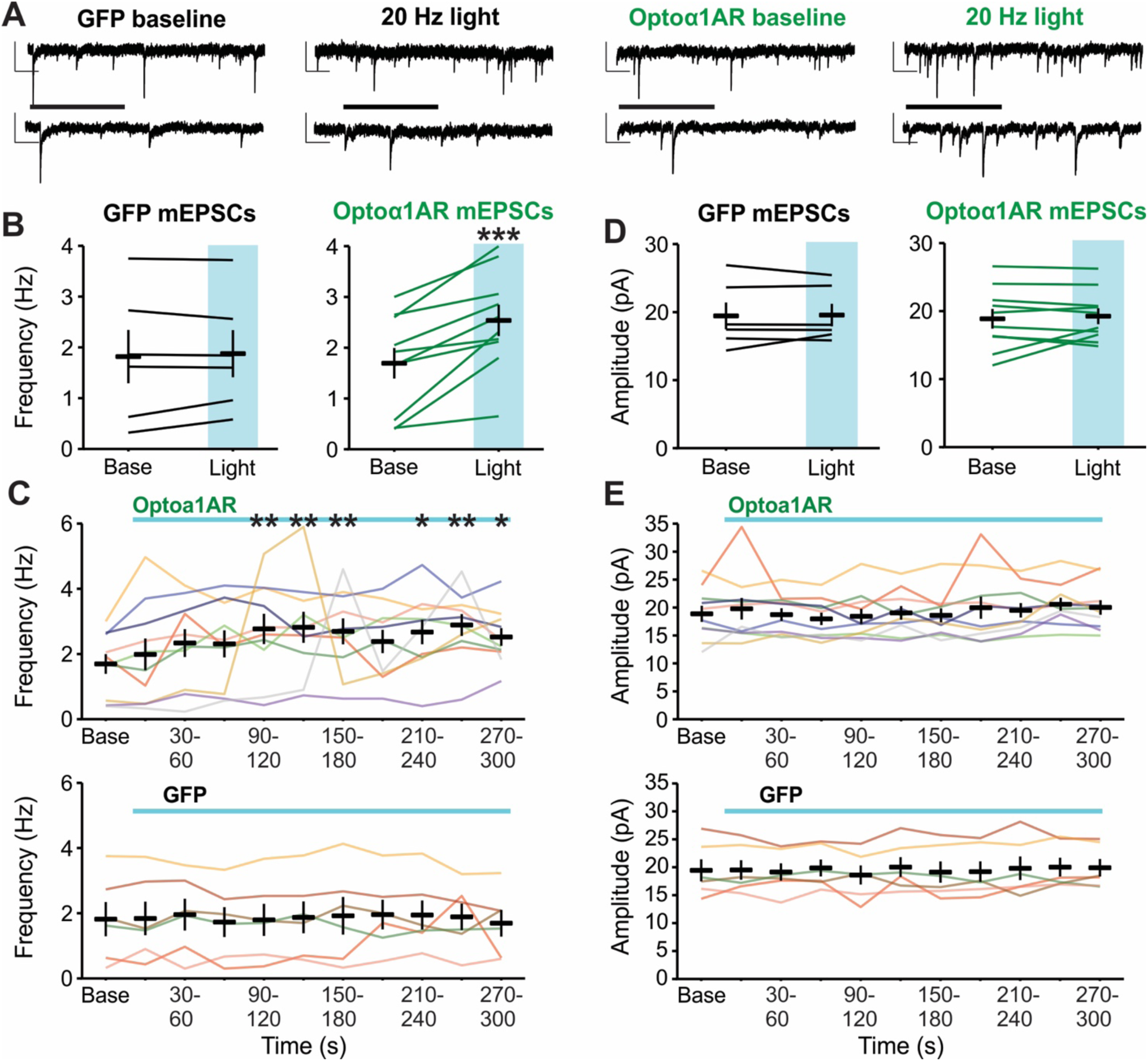
High-frequency (20 Hz) stimulation of astrocytic Optoα1AR increases mEPSC frequency. (A) Representative mEPSC traces from individual pyramidal cells before (left) and during (right) 20 Hz stimulation of GFP (left two traces) or Optoα1AR (right two traces) in astrocytes. Insets represent expanded sections from underlined portion of trace. Scale bars: 20 pA, 500 ms; inset: 20 pA, 100 ms. (B) Mean ± SEM of mEPSC frequency before (Base) and during the full duration of 20 Hz stimulation (Light) of GFP (left) or Optoα1AR (right) in astrocytes. Lines represent individual cells. (C) Mean ± SEM of mEPSC frequency across time in the Optoα1AR (top) and GFP (bottom) groups during 20 Hz stimulation. Lines represent individual cells. Note that lines are used to enable identification of the same cells across the time points; increased slope at end of baseline period does not represent an increase in frequency prior to light delivery. (D) Mean ± SEM of mEPSC amplitude before (Base) and during the full duration of 20 Hz stimulation (Light) of GFP (left) or Optoα1AR (right) in astrocytes. Lines represent individual cells. (E) Mean ± SEM of mEPSC amplitude across time in the Optoα1AR (top) and GFP (bottom) groups during 20 Hz stimulation. Lines represent individual cells. Note that lines are used to enable identification of the same cells across the time points; increased slope at end of baseline period does not represent an increase in frequency prior to light delivery. *, **, ***, p<0.5, p<0.01, p<0.001 vs. baseline value

### Low-frequency (0.5 Hz) optogenetic stimulation of astrocytic ChR2, but not Optoα1AR, modulates hippocampal synaptic inhibition

Responses to optogenetic stimulation are shaped by a combination of the properties of the light stimulation, the kinetics of opsin activation, and the cell expressing the opsin. The Optoα1AR is coupled to a GPCR signaling cascade, which has slower kinetics of activation and deactivation than ionotropic opsins and thus may have a broader range of effective stimulation parameters. Therefore, we tested whether an alternative stimulation paradigm of longer pulses delivered at a lower frequency for a shorter period of time would elicit similar results as the 20 Hz stimulation. Recordings of sIPSCs or sEPSCs were made in CA1 pyramidal cells as before, and slices containing astrocytes expressing either Optoα1AR or control GFP were exposed to 0.5 Hz blue light stimulation at successive 1, 5, or 10 mW intensities for 90 s per intensity, with 2 min between stimulations (**Figure 5A**). Following 0.5 Hz stimulation of slices in the Optoα1AR group, there was no change in sIPSC frequency compared to baseline at any stimulation intensity or timepoint (**Figure 5B and 5C**, **middle**) (n=10 cells from 6 mice). Additionally, there was no difference in sIPSC frequency in response to light stimulation between the Optoα1AR and GFP groups (**Figure 5B and 5C**, **top**) (GFP n=10 cells from 4 mice).

**Figure 5.**
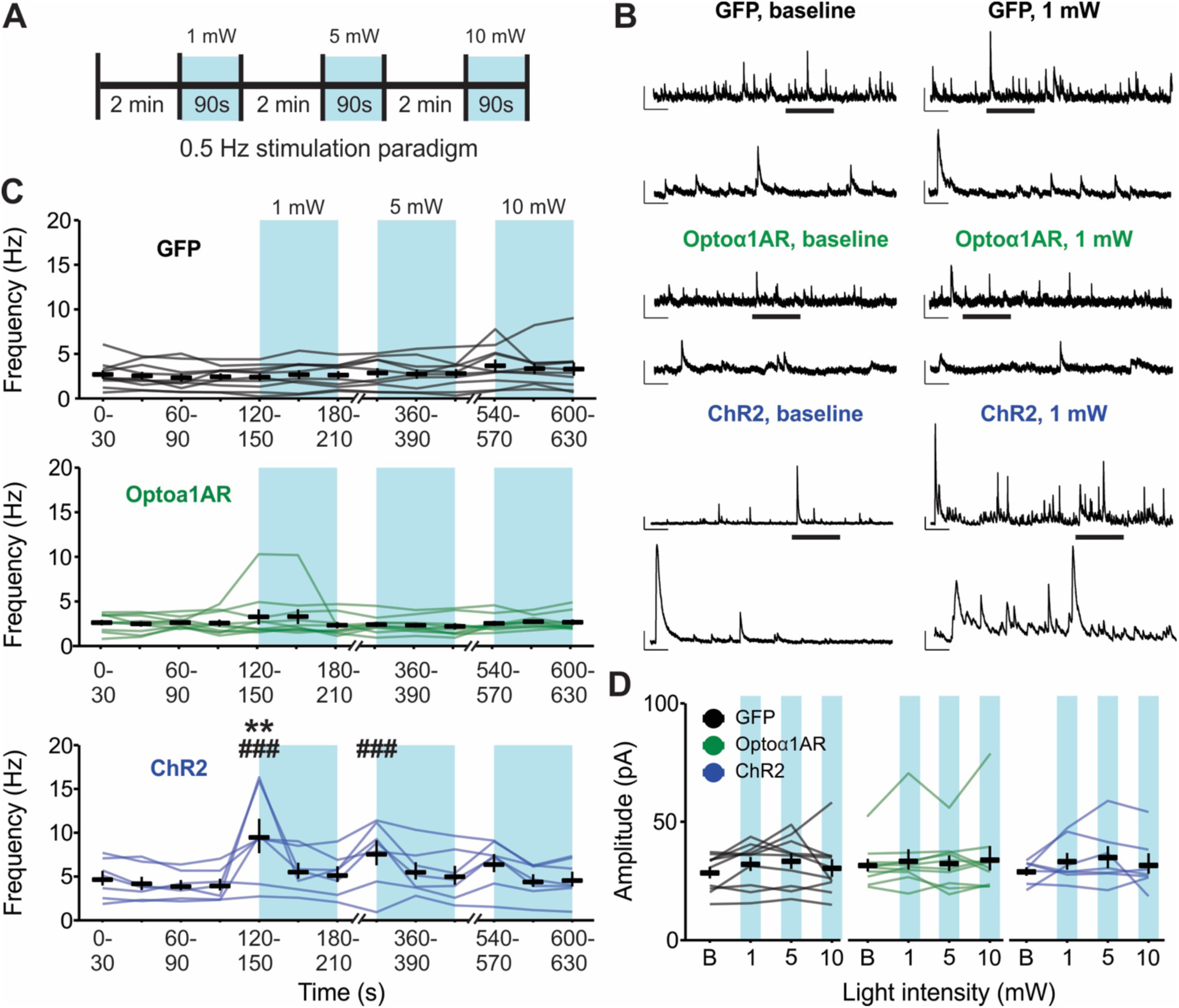
Low-frequency (0.5 Hz) blue light stimulation of astrocytes expressing ChR2, but not Optoα1AR, increases sIPSC frequency. (A) Schematic representation of 0.5 Hz light stimulation paradigm. (B) Representative sIPSC traces from individual pyramidal cells before (left) and during (right) 0.5 Hz blue light stimulation delivered to slices expressing GFP (top), Optoα1AR (middle), or ChR2 (bottom) in astrocytes. Insets represent expanded sections from underlined portion of trace. Scale bars – GFP and Optoα1AR: 50 pA, 1 s; inset: 50 pA, 200 ms; ChR2: 100 pA, 1s; inset: 100 pA, 200 ms. (C) Mean ± SEM of sIPSC frequency across time of GFP (top), Optoα1AR (middle), and ChR2 (bottom) groups during 0.5 Hz stimulation. Lines represent individual cells. Note that lines are used to enable identification of the same cells across the time points; increased slope at end of baseline period does not represent an increase in frequency prior to light delivery. (D) Mean ± SEM of sIPSC amplitude in GFP (left), Optoα1AR (middle), and ChR2 (right) groups across stimulation intensities. Lines represent individual cells. B = baseline. **, p<0.01 vs. baseline values. ###, p<0.001 vs. GFP control group.

As a positive control to ensure that the 0.5 Hz stimulation of astrocytes is capable of driving alterations in hippocampal synaptic transmission, additional cells were recorded in slices with astrocytes expressing the nonspecific cation channel ChR2. The ChR2 group displayed increased IPSC frequency following 1 mW-intensity stimulation compared to baseline (p=0.002) (n=7 cells from 4 mice, 4 of 7 cells responding with >50% frequency increase), as well as compared to the GFP group at 1 mW and 5 mW (p<0.001, respectively) (**Figure 5B and 5C**, **bottom**), confirming that 0.5 Hz light delivery to astrocytes expressing ChR2 can modulate hippocampal synaptic transmission. Notably, the strength of this ChR2-mediated effect diminished over time despite an increase in stimulation intensity; there was no significant difference compared to baseline at 5 mW intensity, nor compared to either baseline or to the GFP group at 10 mW stimulation, indicating a potent rundown of the effect within minutes that increased light intensity did not overcome. Furthermore, no differences were detected in sIPSC amplitude in any experimental group at any stimulation intensity (F=0.76, p=0.55) (**Figure 5D**).

Together, these data indicate that 0.5 Hz optogenetic stimulation of the nonspecific cation channel ChR2 in astrocytes modulates sIPSC frequency in CA1 hippocampal pyramidal cells, but the same stimulation of Optoα1AR to trigger the G_q_ signaling cascade in astrocytes is not effective.

### Low-frequency (0.5 Hz) optogenetic stimulation of astrocytic ChR2, but not Optoα1AR, modulates hippocampal glutamatergic transmission

To determine if low-frequency 0.5 Hz stimulation modulates glutamatergic transmission in the hippocampus, sEPSCs were recorded from CA1 pyramidal cells (**Figure 6A**). Similar to the data for sIPSCs, 0.5 Hz stimulation of slices in the Optoα1AR group did not result in an overall difference in sEPSC frequency compared to baseline at any stimulation intensity (**Figure 6B, middle**) (n=10 cells from 5 mice). In addition, no differences in sEPSC frequency were detected between the Optoα1AR and GFP groups at any stimulation intensity timepoint (**Figure 6B, top**) (GFP n=10 cells from 4 mice).

**Figure 6.**
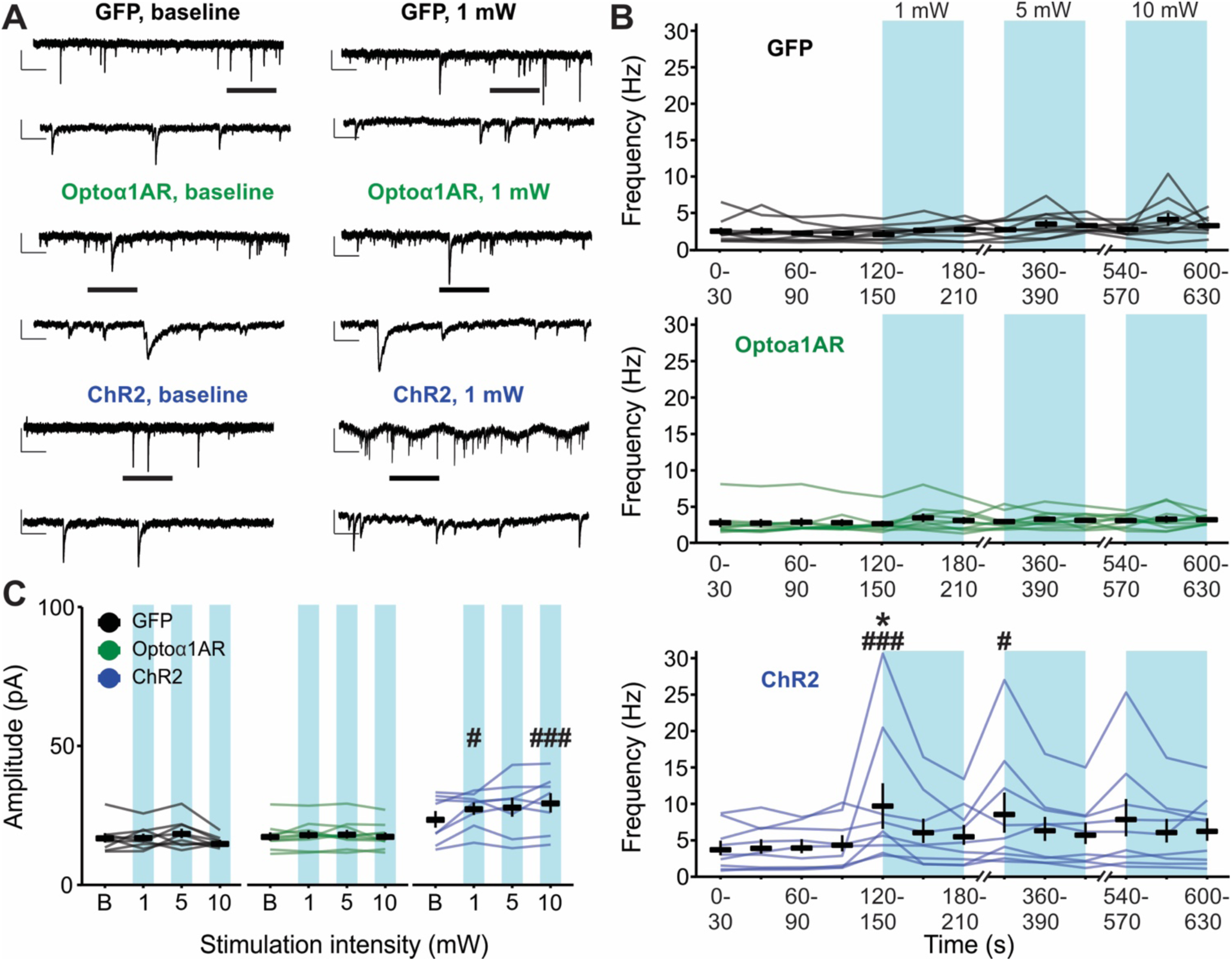
Low-frequency (0.5 Hz) stimulation of astrocytes expressing ChR2, but not Optoα1AR, increases sEPSC frequency and amplitude. (A) Representative sEPSC traces from individual pyramidal cells before (left) and during (right) 0.5 Hz blue light stimulation delivered to slices expressing GFP (top), Optoα1AR (middle), or ChR2 (bottom) in astrocytes. Insets represent expanded sections from underlined portion of trace. Scale bars – GFP and Optoα1AR: 20 pA, 1 s; inset: 20 pA, 200 ms; ChR2: 50 pA, 1s; inset: 50 pA, 200 ms. (B) Mean ± SEM of sEPSC frequency across time of GFP (top), Optoα1AR (middle), and ChR2 (bottom) groups during 0.5 Hz stimulation. Lines represent individual cells. (C) Mean ± SEM of sEPSC amplitude of GFP (left), Optoα1AR (middle), and ChR2 (right) groups across stimulation intensities. Lines represent individual cells. B = baseline. *, p<0.05 vs. baseline values. #, ###, p<0.05, p<0.001 vs. GFP control group.

However, as with sIPSCs, the ChR2 group displayed an increase in sEPSC frequency following 1 mW stimulation as compared to baseline (p=0.04), and compared to GFP controls at both 1 mW (p<0.001) and 5 mW intensities (p=0.02) (n=9 cells from 4 mice, 5 of 9 cells responding with >50% frequency increase) (**Figure 6B, bottom**). This effect also diminished over time, with no difference in response to 10 mW stimulation compared to baseline or compared to control GFP. In contrast to the findings for sIPSCs, however, analysis of sEPSC amplitude with 0.5 Hz stimulation yielded significant effects at 1 mW (p=0.03) and 10 mW (p<0.001) intensities compared to GFP controls (**Figure 6C**). These data indicate that low-frequency 0.5 Hz blue light stimulation does not modulate basal excitatory synaptic transmission in CA1 pyramidal cells when Optoα1AR is expressed in astrocytes, but in the case of astrocytic ChR2, this same stimulation produces increased sEPSC frequency and amplitude.

In 5 of 9 cells, 0.5 Hz stimulation of ChR2 in astrocytes when recording sEPSCs also resulted in the development of an inward tonic current that did not appear when recording sIPSCs. Furthermore, this ChR2-mediated tonic current displayed two unique temporal profiles: a slow tonic current that peaked in amplitude during the first 30 s of 0.5 Hz light exposure (**Figure 7A, top**); and an acute tonic current that fluctuated with the on/off stimulation of the light (**Figure 7A, middle and bottom**). The amplitude of the slow tonic current ranged from 31.98-191.71 pA, with a mean time to peak amplitude of 15.09 s following 1 mW 0.5 Hz stimulation (**Figure 7B**). The acute tonic current was largest during the initial 10 s of 0.5 Hz light stimulation (mean 43.61 ± 20.81 pA), but recovered to 47.54% of max amplitude within 30 s of stimulation (mean 22.88 ± 6.96 pA) (**Figure 7C**). Additionally, a strong linear correlation was observed between the amplitude of the slow tonic current and phasic sEPSC frequency, as cells with the highest phasic sEPSC frequency displayed the greatest amplitude of the slow tonic current (r=0.99, p<0.0001) (**Figure 7D**). Note that this tonic current was not observed in Optoα1AR or GFP groups.

**Figure 7.**
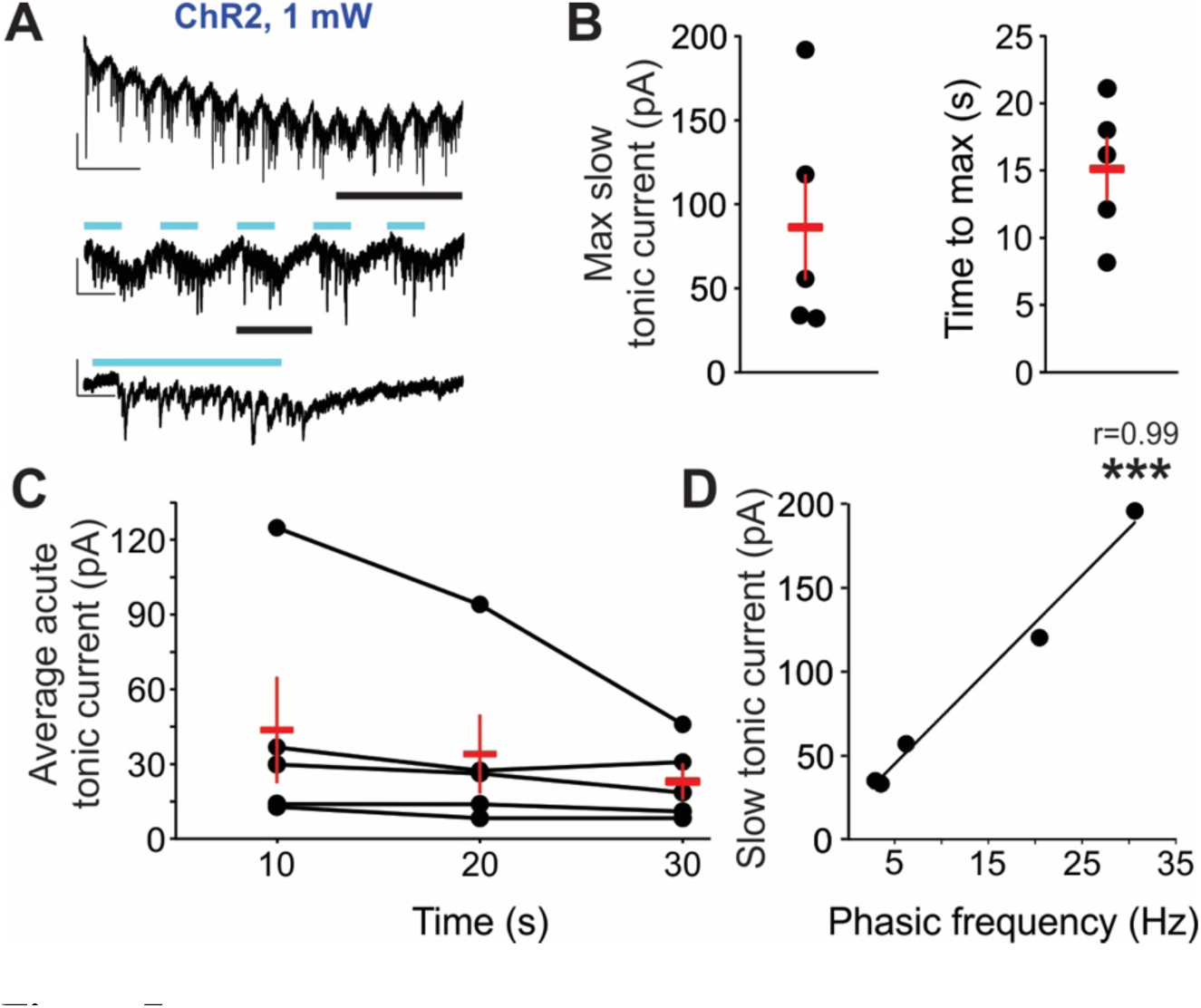
Low-frequency (0.5 Hz) stimulation of astrocytes expressing ChR2 induces both slow and acute tonic inward currents. (A) Representative sEPSC trace from an individual pyramidal cell during 0.5 Hz blue light stimulation delivered to slices expressing astrocytic ChR2. Insets represent expanded sections from underlined portion of trace. Blue lines indicate times of light stimulation. Scale bars: top, 50 pA, 5 s; middle: 50 pA, 1 s; bottom: 50 pA, 200 ms. (B) Mean ± SEM (red markers) of the maximum amplitude of the slow tonic current (left) and time to max amplitude of slow tonic current (right). Dots represent individual cells (5 cells in which tonic currents were elicited, out of 9 recorded). (C) Mean ± SEM (red markers) of the average acute tonic current during first 30 s of blue light exposure. Dots and connected lines represent individual cells. (D) Slow tonic current amplitude plotted in relation to the frequency of phasic sEPSCs in the same cells. Dots represent individual cells; linear regression line of best fit indicates strong correlation. *** p<0.0001, Pearson’s correlation.

## Discussion

In these studies, we tested whether stimulation of the G_q_-coupled opsin, Optoα1AR, can modulate basal synaptic transmission in hippocampal CA1 when expressed in astrocytes. The results indicate that optical stimulation of astrocytic Optoα1AR is capable of modulating synaptic transmission, including basal GABAergic inhibition, in hippocampus. Furthermore, these effects are stimulation-specific; a high-frequency (20 Hz) stimulation of Optoα1AR in astrocytes effectively modulated hippocampal transmission, whereas a low-frequency 0.5 Hz stimulation of Optoα1AR had no effect on modulating IPSCs or EPSCs. The lack of response to 0.5 Hz stimulation was in contrast to large changes seen with low-frequency stimulation of astrocytes expressing the nonspecific cation channel ChR2. 20 Hz stimulation of astrocytic Optoα1AR led to increases in the frequency of sIPSCs, mIPSCs, and mEPSCs, and the increase in mIPSC frequency displayed a trend for reversibility within 20 min. Altogether, these studies provide evidence for stimulation-specific effects of astrocytic Optoα1AR in modulating both GABAergic and glutamatergic transmission in hippocampal CA1. The AAV-GFAP- Optoα1AR vector presented here may thus be a useful tool in probing astrocytic modulation of local synaptic transmission.

Currently, the G_q_-coupled DREADD, hM3Dq^37^, is the most commonly used tool for targeted activation of astrocytes due to its ease of use and physiological relevance when inserted into astrocytes and activated by clozapine-N-oxide (CNO) or other similar ligand. The Optoα1AR vector employed here may be used as an alternative to hM3Dq for astrocyte stimulation by activating the same endogenous pathway with improved temporal specificity. As CNO is typically administered over an extended period of time, and the effects of hM3Dq activation can be long-lasting, one must consider the temporal distinction between experiments that activate astrocytes using DREADDs versus optogenetic tools. The present study reinforces that the frequency and duration of optical stimulation paradigms should be carefully considered when using optogenetic constructs to interrogate roles for astrocytes in modulating synaptic transmission. For example, in a prior study, sustained optical stimulation of CA1 astrocytes expressing G_q_-coupled melanopsin at 7 mW for 1 s or 3 s was insufficient to modulate sEPSCs. However, stimulation lengths of 5 s and 10 s led to transient increases in sEPSC amplitude, and stimulation lengths of 20 s and 60 s induced sustained increases in sEPSC amplitude^9^. The effects on excitatory transmission observed here in response to 20 Hz, 5 mW stimulation of Optoα1AR are in line with these previously described effects of astrocyte-expressed melanopsin and extend the effects of activation of G_q_ signaling in astrocytes to modulation of synaptic inhibition as well. In the present studies, however, 0.5 Hz stimulation (repeated 1-s light pulses delivered over 90 s) of astrocytic Optoα1AR at intensities up to 10 mW did not alter EPSC frequency or amplitude, suggesting that the efficacy of activating Optoα1AR in yielding changes in nearby synaptic transmission is stimulation-sensitive.

The 0.5 Hz stimulation paradigm was designed in an attempt to mimic the relatively slow activation kinetics of astrocytic GPCRs, as well as the temporal dynamics of astrocytic Ca^2+^ signaling, which are generally seconds in length^38, 39^. It should be noted, however, that recent work has identified Ca^2+^ signals in astrocyte microdomains with ms timescales *in vivo*^40^. By contrast, the 20 Hz stimulation was chosen to closely mimic light delivery used in previous *in vivo* experiments testing a similar Optoα1AR opsin expressed in astrocytes^8^. For the 0.5 Hz stimulation, however, it is possible that the escalating-intensity stimulation paradigm used in the present studies masked a potential effect of the higher light intensities. In addition, it is important to note that the stimulation paradigms used in these experiments are just two of many possible options, and the two stimulation paradigms used here differed in the total time of stimulation and whether or not there was an escalation of the stimulation intensity.

Therefore, it is possible that the temporal frequency of the stimulation intensity is not the sole cause for the differences seen in astrocytic Optoα1AR modulation of hippocampal transmission. The present studies were also performed at room temperature to reduce the frequency of synaptic currents and more readily enable acquisition of unitary PSCs, and temperature is known to impact GPCR kinetics. Note, however, that the 20 Hz stimulation paradigm, which demonstrated efficacy at room temperature in the present experiments, was adapted from previous *in vivo* work using a similar Optoα1AR in astrocytes^8^.

The present study reports modulation of basal synaptic inhibition by astrocytes using an optogenetic tool designed to harness physiologically relevant endogenous pathways in astrocytes. High-frequency stimulation of Optoα1AR in astrocytes resulted in elevated mIPSC frequency, which suggests a potential Ca^2+^-dependent release of a signaling molecule from astrocytes that acts upon presynaptic terminals of interneurons, subsequently increasing the probability of GABA release at inhibitory synapses. As mIPSC amplitude was not changed, quantal size does not appear to have been affected.

Purinergic signaling is potential mediator of this response; astrocytes release ATP/adenosine in a Ca^2+^- dependent manner, and ATP/adenosine has been shown to modulate both basal excitatory neurotransmission^15, 41^ as well as heterosynaptic depression^14, 42, 43^ in the hippocampus. Purinergic signaling can also modulate inhibitory transmission; for example, endogenous P2Y_1_ receptor activation in hippocampal interneurons leads to increased interneuron excitability, subsequently increasing feed- forward inhibition onto CA1 pyramidal cells. Both anatomical and functional evidence (a high density of astrocyte processes surrounding hippocampal interneurons and enhanced intracellular Ca^2+^ transients, respectively) suggests that these effects are driven by astrocyte-derived ATP^44^. Additionally, astrocytic release of ATP/adenosine upregulates synaptic inhibition from somatostatin interneurons onto pyramidal cells in the hippocampus^20^. Furthermore, G_q_ pathway stimulation in astrocytes in the central amygdala leads to Ca^2+^-dependent release of ATP/adenosine and an enhancement of inhibition through activation of neuronal A2_A_ receptors^24^. These effects were deemed to be mediated presynaptically, in line with the

increase in mIPSC frequency seen in the present studies. It is important to note that all recordings in this study were made in the absence of synaptic blockers; EPSCs and IPSCs were biophysically isolated only in the recorded cell. Though recordings made from slices in the absence of synaptic blockers may be more representative of *in vivo* conditions, the recordings represent the net result of both excitatory and inhibitory network activity, making the underlying mechanisms difficult to pinpoint. In addition, other potential mechanisms may be considered. For example, astrocytes provide metabolic support to neurons, and disruption of metabolic processes such as glycolysis and oxidative phosphorylation can alter presynaptic vesicular release^45^. Future studies should also consider the potential impact that optical tools may have on the important metabolic roles of astrocytes.

Low-frequency stimulation of ChR2 in astrocytes not only altered phasic synaptic excitation but produced a tonic inward current as well. Of note, a similar slow inward current with superimposed sEPSCs has also been shown after ChR2 activation in astrocytes in the striatum^31^. Mechanistically, the tight correlation between tonic current amplitude and sEPSC frequency in the presence of light stimulation suggests that stimulation of ChR2 in astrocytes causes the release of glutamate from astrocytic stores, flooding synaptic clefts with glutamate and increasing synaptic EPSC frequency at least in part via increased binding events onto postsynaptic ionotropic glutamate receptors. Subsequent glutamate spillover could then drive the tonic inward current, possibly via activation of extrasynaptic NMDA receptors^46–48^, which can be activated despite a hyperpolarized holding potential and the presence of Mg^2+^ in the extracellular solution^47^. However, stimulation of astrocytic ChR2 can also increase local neuronal excitability independently of astrocytic glutamate release. For example, ChR2 stimulation in striatal astrocytes leads to local neuronal depolarization attributed solely to transient increases in extracellular K^+30^.

The present study used a novel variant of Optoα1AR^33^ packaged in AAV8 and driven by a GFAP promoter for astrocyte-specificity. Optoα1AR canonically drives Ca^2+^ elevations via activation of endogenous G_q_, leading to the release of Ca^2+^ from IP_3_-dependent intracellular stores, as previously demonstrated both in HEK cells ^33^ and cultured astrocytes^34^. Therefore, although it is unlikely that the present results are due to a Ca^2+^-independent mechanism triggered in the astrocytes, we cannot discount this possibility. One potential explanation for the stimulation-specific modulation of hippocampal synaptic transmission seen here is that the 0.5 Hz stimulation of Optoα1AR may be insufficient to drive consistent increases in astrocytic intracellular Ca^2+^ levels as compared to the high-frequency 20 Hz stimulation, thus failing to trigger Ca^2+^-dependent release of signaling molecules. Future studies using the GFAP-Optoα1AR construct could compare the properties of astrocytic Ca^2+^ elevations following exposure to varying stimulation paradigms.

In summary, the goal of these experiments was to evaluate the efficacy of activating Optoα1AR in astrocytes in modulating basal synaptic transmission. The present data indicate that high-frequency blue light activation of Optoα1AR in astrocytes can effectively modulate basal synaptic inhibition as well as excitation in the hippocampus. By contrast, low-frequency 0.5 Hz stimulation of astrocytic Optoα1AR did not affect glutamatergic or GABAergic transmission, although this stimulation produced robust responses when ChR2 was expressed in astrocytes. Overall, this work suggests that activation of AAV-transduced Optoα1AR in astrocytes can effectively alter local synaptic transmission, indicating that this vector should be a useful tool in studies of astrocyte-neuron interactions.

## Methods

### Animals

All animal procedures were approved by the Institutional Animal Care and Use Committee of the University of Illinois at Urbana-Champaign (protocols 17161 and 20110). Male and female C57BL/6J mice were either bred on site or obtained from the Jackson Laboratory at 6-8 weeks of age. Mice were group-housed (up to five mice per cage) in a 14/10 h light:dark cycle with food and water available *ad libitum*.

### Viral vectors

The pAAV-GFAP-OptoA1-eYFP plasmid was constructed by replacing the CaMKIIa promoter in pAAV-CaMKIIa-OptoA1-eYFP by a 2.2 Kb GFAP promoter and verified by Sanger sequencing. The map and sequence information are available at: web.stanford.edu/group/dlab/optogenetics/sequence_info.html#optoxr. AAV-8 (Y733F), referred to here as AAV8, was produced by the Stanford Neuroscience Gene Vector and Virus Core. In brief, AAV8- GFAP-OptoA1-eYFP was produced by standard triple transfection of AAV 293 cells (Agilent). At 72 h post-transfection, the cells were collected and lysed by a freeze-thaw procedure. Viral particles were then purified by an iodixanol step-gradient ultracentrifugation method. The iodixanol was diluted and the AAV was concentrated using a 100-kDa molecular mass–cutoff ultrafiltration device. Genomic titer was determined by quantitative PCR. The virus was tested in cultured neurons for expected expression patterns prior to use *in vivo*. AAV8-GFAP-hChR2(H134R)-eYFP and AAV8-GFAP-eGFP were obtained from the UNC Vector Core. All vectors were diluted in 0.9% sterile saline to a final titer of 1#x00D7;10^12^ for injections; all dilutions were performed immediately prior to injection.

### Stereotaxic virus injections

Stereotaxic injections were performed in mice aged postnatal day (P)42 to P90. Animals were anesthetized using 2-3% oxygen-vaporized isoflurane anesthesia (Clipper Distributing Company) and were placed in a stereotactic apparatus (Kopf Instruments). Carprofen (5 mg/kg, Zoetis) was administered subcutaneously at the beginning of surgery for analgesia. Viral vectors were loaded into a 10-μl Nanofil syringe with a 33-gauge needle, and injections were carried out using a Micro4 injection pump controller (World Precision Instruments). Viruses were bilaterally injected (1 μl per site) into dorsal hippocampal CA1 (coordinates: 1.8 mm posterior and 1.3 mm lateral to bregma; 1.3 mm ventral to the cortical surface) at a rate of 0.12 μl/min. After each injection, the syringe was left in place for 3-5 min to allow for diffusion of the viral vector and minimize reflux along the injection track. Incisions were closed using Perma-Hand silk sutures (Ethicon). Following surgery completion, 2.5% lidocaine + 2.5% prilocaine cream (Hi-Tech Pharmacal) and Neosporin antibiotic gel (Johnson and Johnson) were applied to the incision site.

### Brain slice preparation

Acute brain slices were prepared at ages P80-P145, with a range of 26-71 days after stereotaxic virus injection. Mice were anesthetized via intraperitoneal injection of pentobarbital (Vortech Pharmaceuticals, 55 mg/kg) as performed previously ^18, 49, 50^ and euthanized by decapitation. Brains were immediately dissected and placed in an ice-cold oxygenated (95% O_2_/5% CO_2_) high-sucrose slicing solution containing (in mM) 254 sucrose, 11 glucose, 2.5 KCl, 1.25 NaH_2_PO_4_, 10 MgSO_4_, 0.5 CaCl_2_, and 26 NaHCO_3_. 300 μm-thick coronal slices through dorsal hippocampus were prepared using a Leica VT1200S vibratome (Leica Biosystems). Slices were hemisected, transferred to a holding chamber, and incubated in an oxygenated artificial cerebrospinal fluid (ACSF) solution containing (in mM) 126 NaCl, 2.5 KCl, 10 glucose, 1.25 NaH_2_PO_4_, 1 MgSO_4_, 2 CaCl_2_, and 26 NaHCO_3_ at ∼298 mOsm. For all experiments, slices were incubated in ACSF for 60 min at 32°C, then moved to room temperature (21- 23°C) for at least 15 min before recording.

### Patch clamp electrophysiology

Slices were placed in a fully submerged recording chamber on the stage of an upright BX51WI microscope (Olympus America) and continuously superfused with oxygenated ACSF at a rate of 2.5 ml/min at room temperature. Recordings were made using a MultiClamp 700B amplifier, Digidata 1550 digitizer, and Clampex 10 software (Molecular Devices). Recording pipettes were prepared from thick-walled borosilicate glass using a P-1000 micropipette puller (Sutter Instruments). For all experiments, access resistance was monitored every 2-5 minutes. Only cells that displayed a low and stable access resistance (R_a_ <20 MΩ; <20% change in R_a_ for the duration of the experiment) were kept for analyses. There were no differences between groups in access resistance, input resistance, or cell capacitance.

For voltage-clamp recordings, pipettes were pulled to have an open-tip resistance of 2-5 MΩ when filled with an internal solution containing (in mM): 130 Cs-gluconate, 8 CsCl, 2 NaCl, 10 HEPES, 4 EGTA, 4 Mg-ATP, 0.3 GTP, adjusted to 290 mOsm and pH 7.3. Individual neurons were selected for their pyramidal shape using differential infrared contrast optics through either a sCMOS camera (OrcaFlash 4.0LT, Hamamatsu) or a Retiga R1 CCD camera (Teledyne Photometrics). Slices were screened for GFP expression by brief epifluorescence illumination at 470 nm, and only neurons with nearby astrocytes displaying fluorescence were subsequently recorded. Excitatory and inhibitory postsynaptic currents (EPSCs/IPSCs) were recorded in the absence of synaptic blockers. EPSCs were recorded at a membrane holding potential (V_m_) of -70 mV, whereas IPSCs were recorded at V_m_ = 0 mV. The efficacy of this recording paradigm in isolating EPSCs and IPSCs, respectively, was confirmed by the abolishment of currents following application of glutamate receptor blockers APV and DNQX with V_m_ = -70 mV, as well as the abolishment of currents following application of the GABA_A_ receptor blocker picrotoxin with V_m_ = 0 mV. Each cell was randomly assigned for recording of either EPSCs or IPSCs. Only one cell was recorded per slice. For miniature PSC (mEPSC/mIPSC) recordings, 0.5 μM tetrodotoxin (TTX, Abcam) was added to the bath ACSF. For optogenetic activation, 473 nm blue laser light (Laserglow Technologies) was delivered through an optical fiber 200 μm in diameter (FT200EMT, Thorlabs) placed directly above stratum radiatum/stratum pyramidale at the surface of the slice. Two stimulation paradigms were used: 1) 20 Hz (45-ms pulses at 5 mW intensity, 5 min in duration, 90% duty cycle; partially adapted from *in vivo* experiments in Adamsky et al., 2018); 2) 0.5 Hz (1-s pulses at successive 1, 5, and 10 mW intensities, 90-s duration per intensity, 50% duty cycle).

### Immunohistochemistry

At 3 weeks after AAV injection, animals were euthanized via intracardiac perfusion with 20 mL of 0.1M PBS, followed by 20 mL of 4% paraformaldehyde (PFA). Brain tissue was then collected, fixed in 4% PFA for 24 h at 4°C, and preserved in 30% sucrose solution with 0.5% sodium azide until sectioning. 40 μm-thick coronal sections were prepared using a freezing microtome (SM 2010R, Leica Biosystems). Hippocampal sections were washed 3 times with 0.01 M PBS for 5 min each at room temperature on a shaker at 140 rpm, then incubated for 1 h in a tris-buffered saline (TBS)-based blocking solution (10% normal donkey serum, 0.1% Triton X-100, 2% bovine serum albumin). Sections were then incubated with primary antibody (anti-NeuN rabbit polyclonal [1:1000, Sigma ABN78]) for 48 h at 4°C on a shaker. Sections were then washed in PBS and incubated with DyLight 594-conjugated goat anti-rabbit secondary antibody (1:1000, Vector Laboratories DI-1594) for 2 h on a shaker at room temperature. Tissue was mounted on charged glass slides and coverslipped using Vectashield Hardset Antifade Mounting Medium with DAPI (Vector Laboratories, H-1500). Image acquisition was performed using a BX43 epifluorescence microscope equipped with a Q-Color 3 camera (Olympus) and QCapture Pro 7 software (Teledyne Photometrics).

### Data analysis and statistics

Postsynaptic currents were analyzed using Stimfit software^51^ or a custom analysis package in IGOR Pro (WaveMetrics)^52^. Passive electrical properties were calculated using Clampfit 10.4 (Molecular Devices). Data from Stimfit/IGOR were transferred to OriginPro 2016 (OriginLab, Northampton, MA, USA) or RStudio for statistical analysis. Normality assumptions were evaluated using Shapiro-Wilk tests or through analysis of QQ plots, in which normality was confirmed when data points were closely aligned with reference lines. Within-group comparisons for the 20 Hz stimulation experiments (before/after light stimulation) were made using paired t-tests. 20 Hz PSC recordings compared within the same cells over multiple timepoints were analyzed using one-way repeated-measures ANOVA (with time as the repeated factor) with Fisher’s LSD post-hoc tests. Comparisons for 0.5 Hz stimulation experiments were made using two-way repeated-measures ANOVA (opsin and stimulation intensity as factors) with Fisher’s post-hoc tests. Pearson’s correlation coefficient (r) was used to measure the linear correlation between tonic current amplitude and EPSC frequency. P<0.05 was considered statistically significant.

## Data availability

The datasets generated and analyzed during the current study are available from the corresponding author upon reasonable request.

## Acknowledgements

This work was supported by a Whitehall Foundation Research Grant, a Brain Research Foundation Fay/Frank Seed Grant, a CURE Epilepsy Taking Flight Award, NIH/NINDS grant R01 NS105825 (C.A.C.-H.), and the Beckman Institute (C.S. and N.M.W.). We thank Leanna Leverton, Lola Lozano, and Jordyn Robare for assistance with mouse colony maintenance.

## Author contributions

C.D.C., C.S., and C.A.C.-H. designed experiments; C.D.C., C.S., N.M.W. performed experiments; C.D.C., C.S., R.J.I., and C.A.C.-H. analyzed data; C.R., R.A.D., and K.D. contributed unpublished reagents or tools; C.D.C. and C.A.C.-H. wrote the paper. All authors read and approved the final manuscript.

## Ethics declaration

The authors have no competing interests to declare.

## References

1. Haydon, P. G. GLIA: listening and talking to the synapse. Nat. Rev. Neurosci. 2, 185–193 (2001).

2. Araque, A. et al. Gliotransmitters travel in time and space. Neuron 81, 728–739 (2014).

3. Bazargani, N. & Attwell, D. Astrocyte calcium signaling: the third wave. Nat. Neurosci. 19, 182– 189 (2016).

4. Agulhon, C. et al. What is the role of astrocyte calcium in neurophysiology? Neuron 59, 932–946 (2008).

5. Araque, A., Parpura, V., Sanzgiri, R. P. & Haydon, P. G. Tripartite synapses: glia, the unacknowledged partner. Trends Neurosci. 22, 208–215 (1999).

6. Perea, G., Navarrete, M. & Araque, A. Tripartite synapses: astrocytes process and control synaptic information. Trends Neurosci. 32, 421–431 (2009).

7. Santello, M., Calì, C. & Bezzi, P. Gliotransmission and the tripartite synapse. in Synaptic Plasticity 307–331 (Springer, 2012).

8. Adamsky, A. et al. Astrocytic Activation Generates De Novo Neuronal Potentiation and Memory Enhancement. Cell 174, 59–71.e14 (2018).

9. Mederos, S. et al. Melanopsin for precise optogenetic activation of astrocyte-neuron networks. Glia 67, 915–934 (2019).

10. Covelo, A. & Araque, A. Neuronal activity determines distinct gliotransmitter release from a single astrocyte. Elife 7, e32237 (2018).

11. Jourdain, P. et al. Glutamate exocytosis from astrocytes controls synaptic strength. Nat. Neurosci. 10, 331–339 (2007).

12. Perea, G. & Araque, A. Astrocytes potentiate transmitter release at single hippocampal synapses. Science 317, 1083–1086 (2007).

13. Navarrete, M. & Araque, A. Endocannabinoids potentiate synaptic transmission through stimulation of astrocytes. Neuron 68, 113–126 (2010).

14. Boddum, K. et al. Astrocytic GABA transporter activity modulates excitatory neurotransmission. Nat. Commun. 7, 13572 (2016).

15. Panatier, A. et al. Astrocytes are endogenous regulators of basal transmission at central synapses. Cell 146, 785–798 (2011).

16. Durkee, C. A. et al. G i/o protein-coupled receptors inhibit neurons but activate astrocytes and stimulate gliotransmission. Glia 67, 1076–1093 (2019).

17. Mederos, S. & Perea, G. GABAergic-astrocyte signaling: A refinement of inhibitory brain networks. Glia 67, 1842–1851 (2019).

18. Christian, C. A. & Huguenard, J. R. Astrocytes potentiate GABAergic transmission in the thalamic reticular nucleus via endozepine signaling. Proc. Natl. Acad. Sci. 110, 20278–20283 (2013).

19. Kang, J., Jiang, L., Goldman, S. A. & Nedergaard, M. Astrocyte-mediated potentiation of inhibitory synaptic transmission. Nat. Neurosci. 1, 683–692 (1998).

20. Matos, M. et al. Astrocytes detect and upregulate transmission at inhibitory synapses of somatostatin interneurons onto pyramidal cells. Nat. Commun. 9, 4254 (2018).

21. Agulhon, C. et al. Modulation of the autonomic nervous system and behaviour by acute glial cell Gq protein-coupled receptor activation in vivo. J. Physiol. 591, 5599–5609 (2013).

22. Yang, L., Qi, Y. & Yang, Y. Astrocytes control food intake by inhibiting AGRP neuron activity via adenosine A1 receptors. Cell Rep. 11, 798–807 (2015).

23. Chen, N. et al. Direct modulation of GFAP-expressing glia in the arcuate nucleus bi-directionally regulates feeding. Elife 5, e18716 (2016).

24. Martin-Fernandez, M. et al. Synapse-specific astrocyte gating of amygdala-related behavior. Nat. Neurosci. 20, 1540–1548 (2017).

25. Gourine, A. V. et al. Astrocytes control breathing through pH-dependent release of ATP. Science 329, 571–575 (2010).

26. Perea, G., Yang, A., Boyden, E. S. & Sur, M. Optogenetic astrocyte activation modulates response selectivity of visual cortex neurons in vivo. Nat. Commun. 5, 3262 (2014).

27. Yamashita, A. et al. Astrocytic activation in the anterior cingulate cortex is critical for sleep disorder under neuropathic pain. Synapse 68, 235–247 (2014).

28. Masamoto, K. et al. Unveiling astrocytic control of cerebral blood flow with optogenetics. Sci. Rep. 5, 11455 (2015).

29. Tan, Z. et al. Glia-derived ATP inversely regulates excitability of pyramidal and CCK-positive neurons. Nat. Commun. 8, 13772 (2017).

30. Octeau, J. C. et al. Transient, Consequential Increases in Extracellular Potassium Ions Accompany Channelrhodopsin2 Excitation. Cell Rep. 27, 2249–2261.e7 (2019).

31. Cui, Q. et al. Blunted mGluR Activation Disinhibits Striatopallidal Transmission in Parkinsonian Mice. Cell Rep. 17, 2431–2444 (2016).

32. Xie, A. X., Petravicz, J. & McCarthy, K. D. Molecular approaches for manipulating astrocytic signaling in vivo. Front. Cell. Neurosci. 9, 144 (2015).

33. Airan, R. D., Thompson, K. R., Fenno, L. E., Bernstein, H. & Deisseroth, K. Temporally precise in vivo control of intracellular signalling. Nature 458, 1025–1029 (2009).

34. Figueiredo, M. et al. Comparative analysis of optogenetic actuators in cultured astrocytes. Cell Calcium 56, 208–214 (2014).

35. Mederos, S. et al. GABAergic signaling to astrocytes in the prefrontal cortex sustains goal-directed behaviors. Nat. Neurosci. 24, 82–92 (2021).

36. Deisseroth, K. Optogenetics: 10 years of microbial opsins in neuroscience. Nat. Neurosci. 18, 1213–1225 (2015).

37. Roth, B. L. DREADDs for Neuroscientists. Neuron 89, 683–694 (2016).

38. Haustein, M. D. et al. Conditions and constraints for astrocyte calcium signaling in the hippocampal mossy fiber pathway. Neuron 82, 413–429 (2014).

39. Srinivasan, R. et al. Ca2+ signaling in astrocytes from Ip3r2-/- mice in brain slices and during startle responses in vivo. Nat. Neurosci. 18, 708–717 (2015).

40. Stobart, J. L. et al. Cortical circuit activity evokes rapid astrocyte calcium signals on a similar timescale to neurons. Neuron 98, 726–735 (2018).

41. Di Castro, M. A. et al. Local Ca2+ detection and modulation of synaptic release by astrocytes. Nat. Neurosci. 14, 1276–1284 (2011).

42. Andersson, M., Blomstrand, F. & Hanse, E. Astrocytes play a critical role in transient heterosynaptic depression in the rat hippocampal CA1 region. J. Physiol. 585, 843–852 (2007).

43. Serrano, A., Haddjeri, N., Lacaille, J.-C. & Robitaille, R. GABAergic network activation of glial cells underlies hippocampal heterosynaptic depression. J. Neurosci. 26, 5370–5382 (2006).

44. Bowser, D. N. & Khakh, B. S. ATP excites interneurons and astrocytes to increase synaptic inhibition in neuronal networks. J. Neurosci. 24, 8606–8620 (2004).

45. Sobieski, C., Fitzpatrick, M. J. & Mennerick, S. J. Differential presynaptic ATP supply for basal and high-demand transmission. J. Neurosci. 37, 1888–1899 (2017).

46. Angulo, M. C., Kozlov, A. S., Charpak, S. & Audinat, E. Glutamate released from glial cells synchronizes neuronal activity in the hippocampus. J. Neurosci. 24, 6920–6927 (2004).

47. Fellin, T. et al. Neuronal synchrony mediated by astrocytic glutamate through activation of extrasynaptic NMDA receptors. Neuron 43, 729–743 (2004).

48. Le Meur, K., Galante, M., Angulo, M. C. & Audinat, E. Tonic activation of NMDA receptors by ambient glutamate of non-synaptic origin in the rat hippocampus. J. Physiol. 580, 373–383 (2007).

49. Courtney, C. D. & Christian, C. A. Subregion-Specific Impacts of Genetic Loss of Diazepam Binding Inhibitor on Synaptic Inhibition in the Murine Hippocampus. Neuroscience 388, 128–138 (2018).

50. Christian, C. A. et al. Endogenous positive allosteric modulation of GABAA receptors by diazepam binding inhibitor. Neuron 78, 1063–1074 (2013).

51. Guzman, S. J., Schlögl, A. & Schmidt-Hieber, C. Stimfit: quantifying electrophysiological data with Python. Front. Neuroinform. 8, 16 (2014).

52. DeFazio, R. A., Elias, C. F. & Moenter, S. M. GABAergic transmission to kisspeptin neurons is differentially regulated by time of day and estradiol in female mice. J. Neurosci. 34, 16296– 16308 (2014).

